# Positive fitness effects help explain the broad range of *Wolbachia* prevalences in natural populations

**DOI:** 10.1101/2022.04.11.487824

**Authors:** Petteri Karisto, Anne Duplouy, Charlotte de Vries, Hanna Kokko

## Abstract

The bacterial endosymbiont *Wolbachia* is best known for its ability to modify its host’s reproduction by inducing cytoplasmic incompatibility (CI) to facilitate its own spread. Classical models predict either near-fixation of costly *Wolbachia* once the symbiont has overcome a threshold frequency (invasion barrier), or *Wolbachia* extinction if the barrier is not overcome. However, natural populations do not all follow this pattern: *Wolbachia* can also be found at low frequencies (below one half) that appear stable over time. *Wolbachia* is known to have pleiotropic fitness effects (beyond CI) on its hosts. Existing models typically focus on the possibility that these are negative. Here we consider the possibility that the symbiont provides direct benefits to infected females (e.g. resistance to pathogens) in addition to CI. We discuss an underappreciated feature of *Wolbachia* dynamics: that CI with additional fitness benefits can produce low-frequency (*<* 1*/*2) stable equilibria. Additionally, without a direct positive fitness effect, any stable equilibrium close to one half will be sensitive to perturbations, which make such equilibria unlikely to sustain in nature. The results hold for both diplodiploid and different haplodiploid versions of CI. We suggest that insect populations showing low-frequency *Wolbachia* infection might host CI-inducing symbiotic strains providing additional (hidden or known) benefits to their hosts, especially when classical explanations (ongoing invasion, source-sink dynamics) have been ruled out.

## Introduction

Maternally inherited endosymbiotic bacteria of genus *Wolbachia* are widespread in arthropods and nematodes (Hilgenboecker et al., 2008; Weinert et al., 2015; Zug and Hammerstein, 2012). Since the discovery of *Wolbachia*-induced cytoplasmic incompatibility (CI, Laven, 1956; Yen and Barr, 1971) – embryonic mortality in the offspring of parents of different infection status – the symbionts have attained considerable empirical as well as theoretical interest. CI is an obvious and dramatic fitness effect, but *Wolbachia*’s effects are often pleiotropic, impacting the fitness of its hosts directly and regardless of the type of mates that the host reproduces with (for example Hoffmann, Turelli, and Harshman, 1990). Assuming that pleiotropy includes direct costs of infection, it becomes at first sight difficult to understand how *Wolbachia* can spread and reach high frequencies. However, theory soon showed that costly *Wolbachia* infections show positive frequency dependence. At low frequencies, costs predominate and *Wolbachia* cannot spread, but if the initial frequency of *Wolbachia* is sufficiently high (above an invasion barrier), then the frequency-dependent success favours the CI-inducing symbiont sufficiently for it to spread (Caspari and Watson, 1959; Fine, 1978). At equilibrium it will be close to fixation, only prevented from fixing by some infections failing to be passed on to offspring (Engelstädter and Telschow, 2009; Hoffmann, Turelli, and Harshman, 1990). This fits in with the general expectation that positive frequency dependence makes it easy to explain either absence or (near) fixation of a trait (Lehtonen and Kokko, 2012).

In many empirical systems, *Wolbachia* are indeed very prevalent in their host populations (Deng et al., 2021; Duplouy and Brattström, 2018; Hoffmann, Turelli, and Harshman, 1990; Jeong et al., 2009) in line with existing theory. However, the recent accumulation of studies on the prevalence and penetrance of *Wolbachia* in diverse host species and populations show a wider range of infection prevalences. Sazama et al. (2019) have reported that infection prevalence of *Wolbachia* is likely to remain below one half in the majority of insect species (Arthofer et al., 2009; Hughes et al., 2011; Sun et al., 2007; Tagami and Miura, 2004), including systems where the spread of *Wolbachia* is potentially ongoing (Duplouy, GD Hurst, et al., 2010). There are also examples of low prevalence remaining reasonably stable over time (Duplouy, Couchoux, et al., 2015). Although some cases of low and apparently stable prevalence might be explicable as a result of migration to sink populations from sources where symbiont prevalence is high (Flor et al., 2007; Telschow et al., 2007), it appears difficult to consider spatial dynamics a sufficient explanation for widespread cases of empirically documented low prevalences.

Here we show that spatial heterogeneity is not needed for models to produce low stable frequencies of CI-inducing symbionts, once one allows the effects of infection on the host to be positive. Studies allowing positive fitness effects of CI-inducing symbionts are rare. However, Zug and Hammerstein (2018) investigated the effect of positive fitness on the invasion dynamics of CI and other types of reproductive parasites into virgin populations and against each other. They demonstrated the key role of relative fitness in determining invasion conditions of the parasites. The effect is often observed as a lower (or absent) invasion threshold frequency. While their focus was mostly on the invasibility of the parasites, we aim to widen the understanding of the resulting stable equilibrium frequencies when positive fitness effects are present.

We first revisit classic CI models (following the notation of Engelstädter and Telschow, 2009) and show that the assumption structure, where infected hosts experience either no or negative fitness effects, cannot equilibrate at infection prevalences below one half (as stated by Turelli, 1994, for a diplodiploid case). We also argue that within the range predicted by the classic model (between one half and one), the lower end of the range is difficult to maintain in the presence of stochastic fluctuations in infection prevalence, as the invasion barrier will in these cases be too close to the stable equilibrium. We then show that when we allow infected hosts to benefit from their infection status, the stable equilibria can exist below one half in a wide range of cases, including situations with and without an invasion barrier. The results of the classical diplodiploid model of CI are quite similar to the results of different haplodiploid models (with an exception that we discuss). Our observations of low frequency stable equilibria (specifically, below one half) are not entirely novel: the analysis of (Zug and Hammerstein, 2018) includes a demonstration of such a situation in a diplodiploid system (their Supplementary Fig. S1), albeit without any written comment as the focus of these authors was elsewhere. Hence, our aim is to strengthen the appreciation of the dynamic consequences of positive fitness effects in the dynamics of CI-inducing symbionts.

## Models and analysis

### Diplodiploid model

Hoffmann, Turelli, and Harshman (1990) presented a seminal model for the spread of CI-inducing symbionts in diploid-diploid species, building upon a slightly different model of Fine (1978). We use the discrete time model of Hoffmann, Turelli, and Harshman (1990) but update it to the notation of Engelstädter and Telschow (2009) (Fig. 1). *Wolbachia*-bearing females have relative fecundity *f* compared to the uninfected baseline (which we set to unity). *Wolbachia* is transmitted maternally to a proportion *t* of eggs. Under rules of CI, an uninfected egg that is fertilized by a *Wolbachia*-modified sperm (produced by an infected male) dies with probability *L*, while *Wolbachia*-infected eggs survive irrespective of the status of the fertilizing sperm (to be precise, this requires sperm to be either from a *Wolbachia*-free father or from a father with the same or a compatible *Wolbachia* strain as the egg; for simplicity we follow earlier studies and consider only one strain in our model).

**Figure 1.**
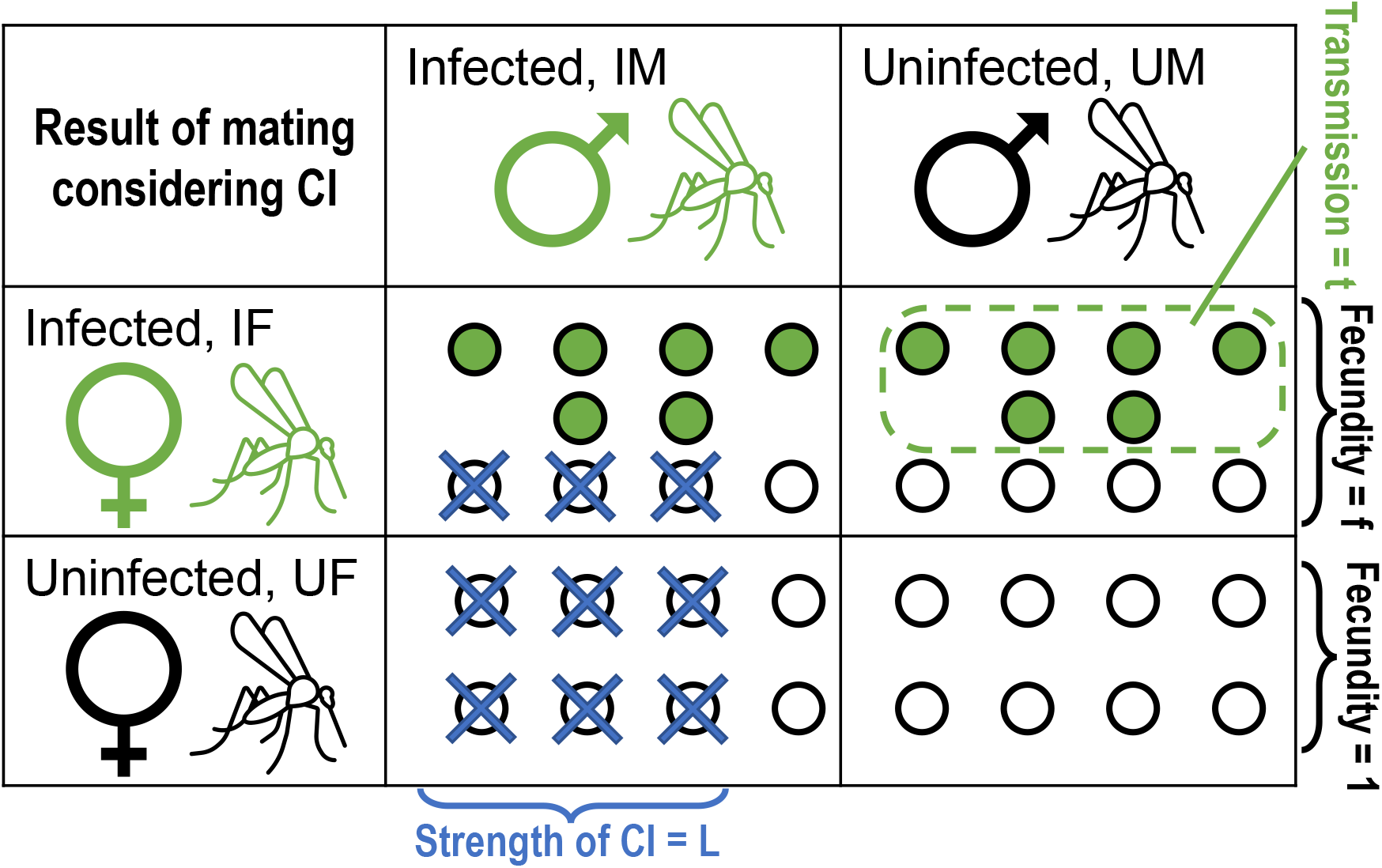
Mating results and key parameters of reproduction when insect population contains *Wolbachia* infected individuals. When an infected male (IM) fertilizes eggs, the *Wolbachia*-modified sperm leads to cytoplasmic incompatibility in all *Wolbachia*-free eggs, regardless of the infection status of their mother (IF, UF). Uninfected males (UM) fertilize all eggs successfully. Note that in the diplodiploid case, the incompatible eggs (that die) could have developed as either males or females. In haplodiploids, where fertilization produces females, the incompatible eggs would all be originally female. In the female-killing type they simply die, while in the masculinization type they develop as males instead.

Our model assumes equal frequency (*p*) of *Wolbachia* in males and females in the current generation. We track four types of matings (the mother and the father can each be uninfected or infected) and their outcome in terms of infected and uninfected offspring production. Only infected mothers can produce infected offspring, and the total of such offspring is *pft* (sperm genotype does not matter for these mothers). The denominator consists of all offspring from all matings. The change from the current generation to the next is

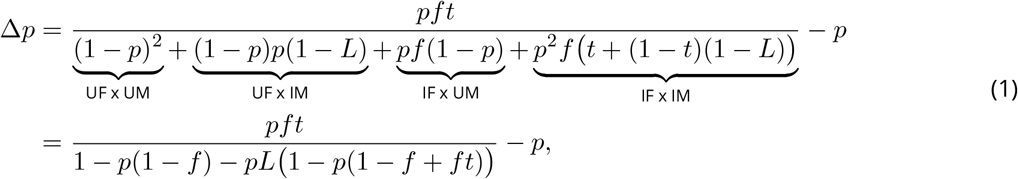

where the first term gives the infection frequency in the next generation. Equation (1) has three equilibria:

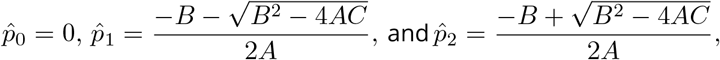

where *A* = *L* (1 − *f* (1 − *t*)), *B* = *f* − 1 − *L* and *C* = 1 − *ft* (Fig. 2a). As shown by Hoffmann, Turelli, and Harshman (1990), when all three equilibria exist, 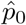 and 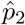 are stable while 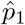 between them is unstable and gives the invasion threshold (in Appendix B we rederive this result extending it to *f >* 1). Turelli (1994, Eq. 4) and Turelli and Hoffmann (1995, Eq. 3) indicated that 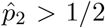, although they did not provide an explicit proof for that. Below, we show what is required for this result to hold.

**Figure 2.**
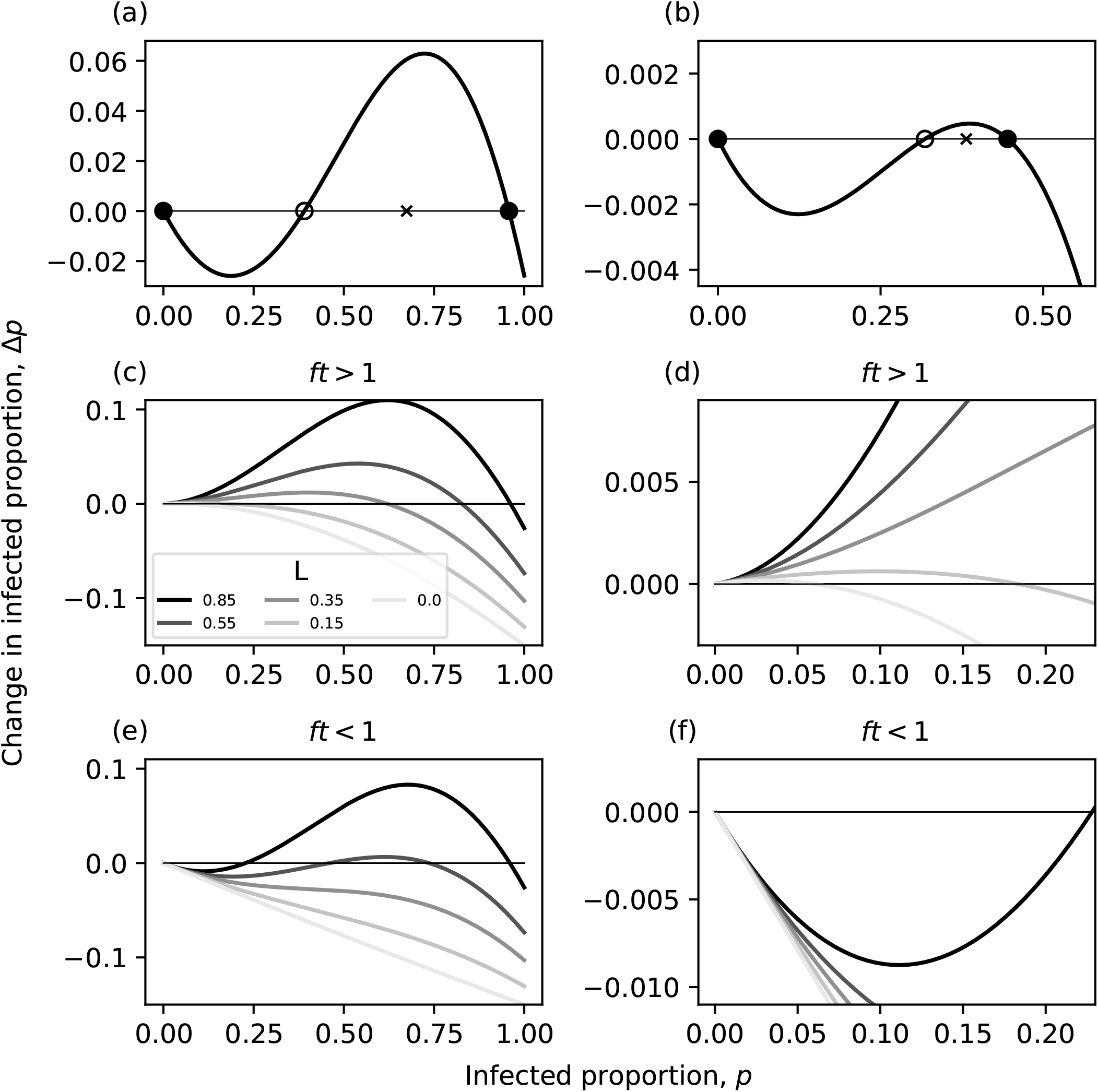
Equilibria of the diplodiploid model (Eq. (1)) shown with plots of Δ*p* as a function of *p*. Dots: stable equilibria, circle: unstable equilibrium, cross: point *r*. a) Classical example with an invasion threshold and high-frequency stable equilibrium. Parameters *f* = 0.85, *t* = 0.85, *L* = 0.85. b) It is possible to have a low stable frequency (*p <* 1*/*2) together with an invasion threshold (*ft* = 0.9588 *<* 1). (*f* = 1.128, *t* = 0.85, *L* = 0.35.) Panels c), d) and e), f) contrast the effect of the strength of CI (*L*) on the invasion dynamics when *ft >* 1 or *ft <* 1. In c),d) *f* = 1.19, *t* = 0.85, and no invasion threshold exists. Low value of *L* leads to low stable infection frequency. In e),f) *f* = 0.99, *t* = 0.85. High *L* shows the classic case, while low *L* predicts extinction of *Wolbachia*.

Before proceeding to understand effects caused by *f >* 1, it is useful to understand why the classical result arises, i.e. that *f* ≤ 1 predicts the absence of low (*<* 1*/*2) and stable frequencies. We outline the reasons both mathematically and biologically. Mathematically, 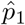 and 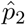 are symmetric around a point *r* = −*B/*(2*A*). Now, for 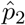 to remain below 1*/*2, it is necessary (but not sufficient) that *r* is below 1*/*2. Solving the inequality *r <* 1*/*2 for *f* with 0 *< t* ≤ 1, 0 *< L* ≤ 1 yields

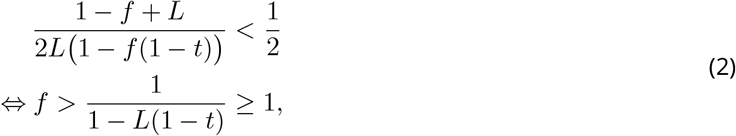

which clearly implies that *r >* 1*/*2 for any *f* ≤ 1. However, the result also suggests that permitting *f >* 1 can change the results, since *r <* 1*/*2 becomes possible and thus possibly also 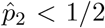. A numerical example shows that this indeed happens with suitable parameter values (Fig. 2b, c, d).

It is worthwhile to consider the robustness of the equilibria against stochastic variation in the environment, that may perturb populations away from their current equilibria. We show that this makes equilibria close to one half unlikely to persist in the long term, within the classically assumed parameter regime where *f* ≤ 1. Consider a situation where *f* ≤ 1 and the stable equilibrium 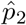 is close to one half. We have a situation 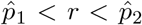, but given the assumption *f* ≤ 1 we also know that *r >* 1*/*2. Since we assumed 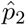 is close to one half, it follows that all points 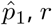 and 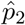 must be close to each other (since *r* is located exactly midway between 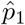 and 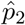). Hence, if the stable equilibrium 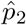 is close to one half, then the unstable equilibrium 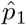 (representing the invasion barrier) is not far below 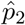. Under such circumstances, random fluctuations in infection prevalence can easily make the population fall stochastically under the invasion barrier, from where it deterministically proceeds towards extinction (unless another stochastic event brings it back up to above the invasion barrier).

Biologically, the above analysis implies that if *Wolbachia* infection has effects that go beyond CI, then these additional fitness differences between infected and uninfected hosts can change the outcome of the interaction significantly. The direction of these differences determine whether *Wolbachia* frequencies can only stabilize at high values (with direct negative fitness effect), or whether the range of prevalence values extends to include low frequencies.

To develop biological intuition for why this is the case, it is instructive to see why the invasion threshold exists in the classical situation (*f* ≤ 1), i.e. why in Fig. 2a there is a switch from poor spread of *Wolbachia* at low frequencies to better success at moderate frequencies. The impact of *Wolbachia* on uninfected eggs is captured by parameter *L*, but the value of *L* becomes irrelevant as a determinant of the dynamics when the infection is rare (very low *p*). A Taylor series expansion of Eq. (1) around *p* = 0 is (*ft*−1)*p*+*f* (1−*f* +*L*)*tp*^2^+ 𝒪 [*p*] ^3^, showing that parameter *L* is associated with second or higher order terms relative to *p*. Hence, when *p* is low, the terms that include *L* become negligible and Δ*p* ≈ (*ft* − 1)*p*.

For the infection to increase in frequency when it is rare, the requirement is that Δ*p >* 0, creating a simple condition: *ft >* 1 means that the ‘leakage’ of offspring into an uninfected state (i.e., only a proportion *t* of offspring of *Wolbachia*-infected mothers inherit the infection) is more than compensated for by the fecundity advantage of *Wolbachia*-infected mothers. Assuming this is the case, the equilibrium 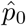 is unstable and *Wolbachia* can spread from rare. Hence there is no invasion threshold when *ft >* 1 (See also Appendix B). Simultaneously, fixation is not possible if we assume *t <* 1; the ‘leakage’ always maintains a supply of uninfected individuals.

Figure 2a shows an example where *ft* is below 1 and spread from rare is consequently not possible. When there are not enough males causing the ‘competing’ egg type to fail (very low *p*), the combination of low fecundity (*f <* 1) and imperfect transmission (*t <* 1) together prevent CI-inducing *Wolbachia* from maintaining its frequency in the population. At higher *p*, we can no longer ignore the consequences of CI, and therefore the effect of the parameter *L*. Now, a high proportion of infected males will elevate the relative success of infected eggs compared to uninfected competitors; the success difference arises from uninfected females’ eggs dying. This explains why in classical examples the CI-inducing symbiont can begin to spread at some threshold value of *p*: the spread is caused by sufficiently many infected males harming the reproduction of uninfected individuals.

The central role of the product *ft* provides important insight. Its value exceeds 1 when *f >* 1*/t*, thus sufficiently high values of *f* can make the invasion barrier disappear (as stated e.g. by Zug and Hammerstein, 2018). However, this can never happen for any *f* ≤ 1, since 1*/t* ≥ 1 for any *t* ∈ (0, 1]. Thus *f >* 1 is a necessary (but not sufficient) requirement for the invasion barrier to disappear. Note that the above holds assuming that the stable polymorphic equilibrium exists but it is possible to lose both the threshold and the stable equilibrium also with *ft <* 1 as seen in Fig. 2. Biologically, situations of *f >* 1*/t >* 1 are possible, once one considers the full range of possibilities for pleiotropy. If females infected by CI-inducing *Wolbachia* are more fecund than uninfected females, and if this advantage is sufficiently large, then the invasion barrier disappears. In this case, one might at first sight expect that this additional advantage would simply make *Wolbachia* reach fixation. This is not the case, however, since the effect of *t <* 1 causing fixation to be impossible still holds.

Instead, the more interesting finding is that *f >* 1 creates conditions where a low frequency of *Wolbachia* can be stable. This was not possible when *f* ≤ 1, because infected males at low prevalence do not cause a large enough survival difference between eggs. But if *f >* 1, infected females can outcompete uninfected ones with very little (assuming *ft <* 1) or no (*ft >* 1) CI-related advantages, and as a whole, the infection can spread at values of *L* that are so low that they would never make *Wolbachia* spread in situations involving *f* ≤ 1 (compare Fig. 2c to Fig. 2e). In other words, the latter condition would not feature an invasion barrier above which positive frequency dependence would be sufficiently strong to bring *Wolbachia* to higher frequencies.

If we now assume that *f >* 1*/t*, we know that the invasion barrier is absent since infected females are more fecund than uninfected ones (as the above implies that *f >* 1). Once the infection spreads, so that infected males are no longer negligibly rare, infected females reap additional benefits (indirectly, by the effect of *L* killing their competitors). The remaining equilibrium in the 0 *< p <* 1 range is stable. Even when *L* = 0, i.e. the effect of males on eggs is switched off and the aforementioned additional benefits are absent, qualitatively similar dynamics arise: the infection frequency increases from rare but cannot fix due to imperfect transmission (also shown in Fig. S1 of Zug and Hammerstein, 2018).

Figures 2c and 2d show examples with *ft >* 1 and five values of *L*. As expected, the *L* = 0 case (lowest curve) exhibits simple dynamics, with the infected strain increasing until imperfect transmission leads to a stable equilibrium which, by solving 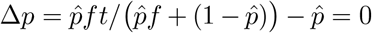, is at 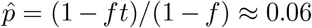 with the given numerical example. Higher values of *L* lead to equilibria with a higher prevalence of *Wolbachia* (Figs. 2 and 3). Choosing equivalent values for *L* in a setting where *ft <* 1 always leads to lower growth (all graphs are lower in Fig. 2e, f compared with the equivalent graphs in Fig. 2c, d). At high *L* this means introducing an invasion barrier, while at low *L* the result is negative growth regardless of the current value of *p*, i.e. extinction of *Wolbachia* (three lowest curves in Fig. 2e).

**Figure 3.**
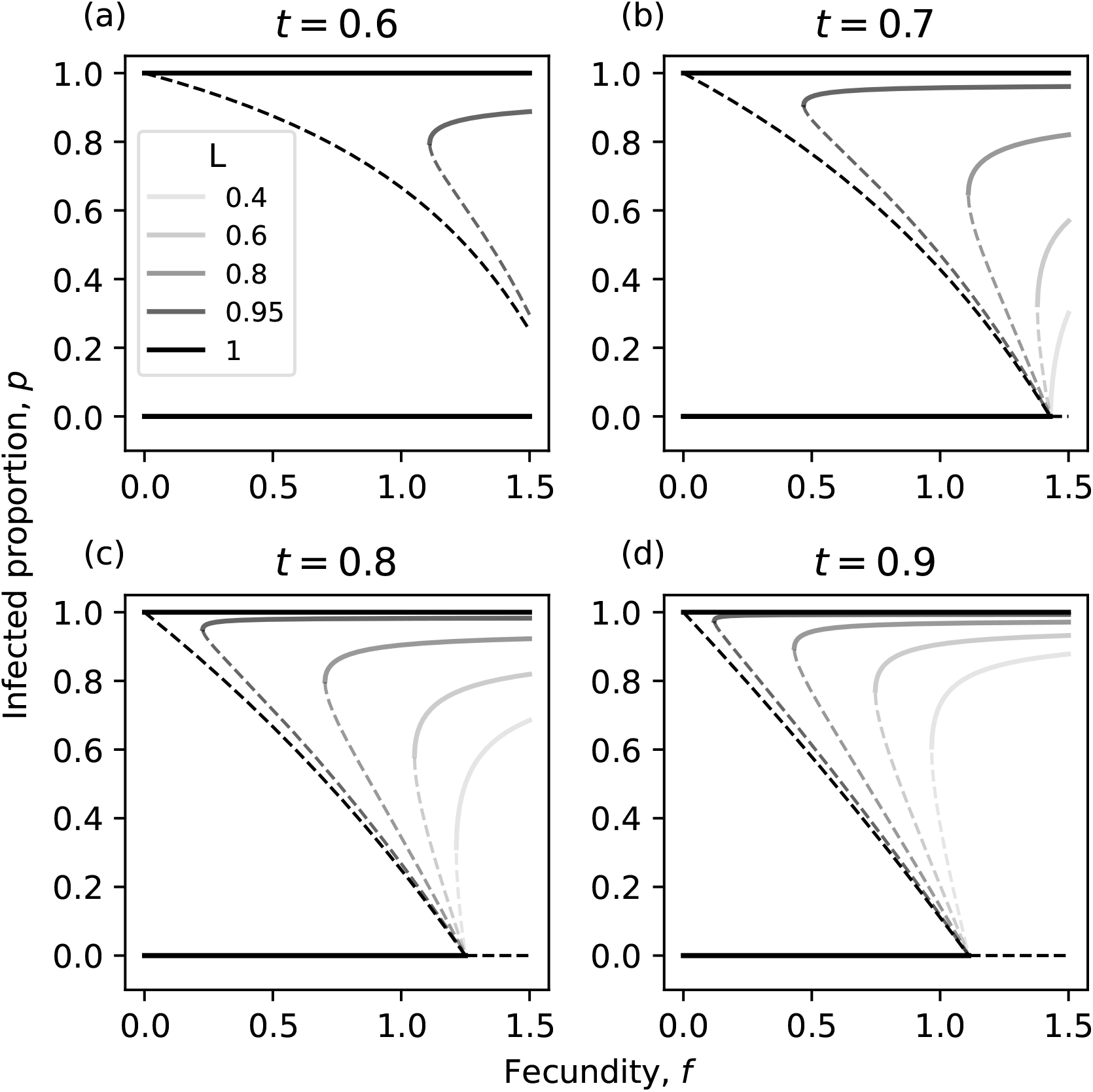
Equilibria of the diplodiploid model (Eq. (1)) as function of *f*, for different values of *t* (panel titles) and *L* (colours of the lines, see legend). Solid lines: stable equilibria. Dashed lines: unstable equilibria.

It is worth clarifying the statement that introducing benefits can allow for a lower equilibrium frequency of *Wolbachia*, as it is counter-intuitive at first sight. The meaning of the statement is not that higher *f* lowers the equilibrium frequency when other parameters are kept constant; this does not happen, instead, higher *f* increases the frequency (Fig. 3). The statement refers to the fact that this very effect (high *f* improves the prospects for *Wolbachia*) can, under parameter settings that are a priori unfavourable to *Wolbachia*, shift a situation where no *Wolbachia* persist to one where some spread is possible, and the system finds its equilibrium at low *p* (the light-coloured curves in Fig. 2c compared to Fig. 2e; Fig. 3). The interesting findings of equilibria with *p <* 1*/*2 arise because *ft >* 1 implies no invasion barrier irrespective of the value of *L*, thus *Wolbachia* can spread from rare even if *L* is meagre, but the modest value of *L* then gives very little ‘boost’ at higher values of *p* (meagre male effects on uninfected eggs when *L* is low).

Although our analysis so far appears to suggest that cases with invasion barrier associate with high equilibrium frequencies, it should be noted that examining the full range of possibilities permitted by *f >* 1 includes cases where an invasion barrier exists, and the stable equilibrium is below one half. An invasion threshold exists if *ft <* 1 (assuming existence of the stable polymorphic equilibrium), while the necessary condition for 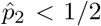 is that *f >* 1. Suitably chosen values for *f* and *t* can fulfil both criteria simultaneously (example: Fig. 2b). As a summary, increasing *f* will increase the value of the stable equilibrium, but at the same time open possibilities for less successful symbionts, manifested as lower values of *L* and *t*, that will equilibrate at lower frequencies. The effect of parameter *f* with different combinations of *t* and *L* is shown in Fig. 3.

It is of interest to see if equilibria with low *p* can only exist at very low values of *L* (mild incompatibility) as in the above examples, or whether other values of other parameters allow low-*p* equilibria to exist even if *L* is relatively high. The latter proves to be the case, and as a whole, assuming *f >* 1 reveals a richer set of possible outcomes than what can occur if *f* ≤ 1 (Fig. 4). The “classical” example with two stable equilibria and an invasion threshold between them (light grey in Fig. 4), as well as extinction of *Wolbachia* (white), can be found whether *f* indicates costs or benefits to being infected, but *f >* 1 additionally permits cases without invasion barriers and with low stable prevalence of *Wolbachia* when low or moderate *L* combines with *ft >* 1 (dark grey areas). Finally, the situation with the stable equilibrium below one half co-occurring with an invasion threshold places conflicting demands on the value of *f* (see above), and while not impossible to fulfil, satisfying them requires rather fine tuned parameter settings with low or intermediate values of *L* and for a narrow range of *t* slightly below 1*/f* (black areas in Fig. 4).

**Figure 4.**
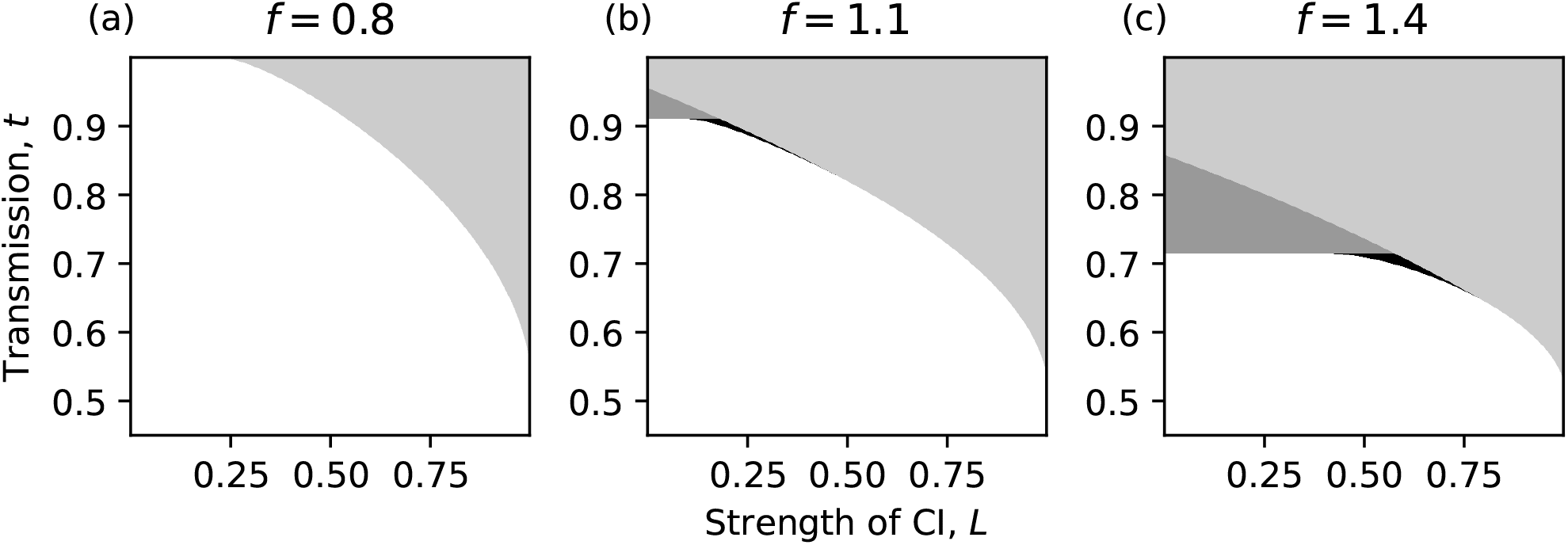
Properties of equilibria of the diplodiploid model shown across parameter space. The parameter regions are categorized as featuring only the trivial equilibrium at *p* = 0 (white), a non-trivial stable equilibrium 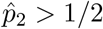 with or without invasion threshold (light grey), and non-trivial stable equilibrium 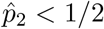 without (dark grey) or with (black) an invasion threshold. Values of parameter *f* (fecundity) are indicated above each panel.

### Haplodiploid models

Thus far, the presented analysis of low stable frequencies of *Wolbachia* assumed diplodiploid sex determination. In this section, we extend these results to two haplodiploid cases, that differ in the effects of an infection on the host. The first case resembles the classic diplodiploid model in that cytoplasmic incompatibility kills fertilized eggs, but since under haplodiploidy all fertilized eggs develop as females, we now switch to calling it the female-killing effect (and note that it is equivalent to the “*Leptopilina* type” of Vavre et al. (2000)). In the second case, incompatibility instead leads to the loss of paternal chromosomes and haploidization of the zygote, which consequently develops as a male (Breeuwer and Werren, 1990). We call this the masculinization effect (Vavre et al. (2000) call it the “*Nasonia* type”, see also Breeuwer and Werren (1990)). Note that both the female-killing effect and the masculinization effect can lead to a male-biased sex ratio, but the underlying mechanisms differ, and we model them separately.

The existence of a low-value stable equilibrium of *Wolbachia* with *f >* 1 also proves true in these two haplodiploid systems. We consider the female-killing effect first. Applying our notation from above to the model of Vavre et al. (2000), the dynamics for *Wolbachia* frequency in females and males, *p*_*F*_ and *p*_*M*_ respectively, become

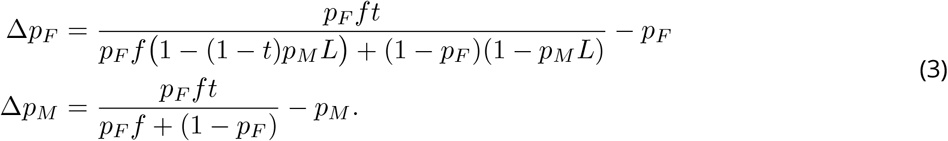

For the masculinization effect, we again adopt the model from Vavre et al. (2000); the dynamics for females and males, respectively, are given as

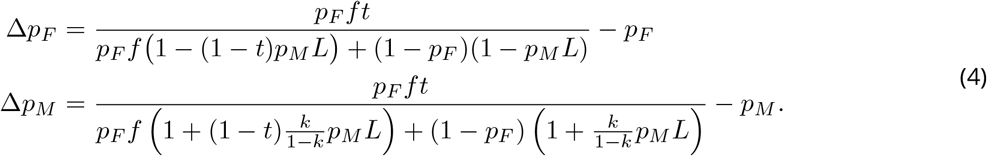

The female equations are identical across the two effects (Eqs. (3) and (4)), as females are lost due to infections impacting fertilized eggs in both cases. However, the male equations differ, because the masculinization effect converts females into males instead of killing them. The magnitude of this additional “male production” depends on the fertilization rate *k*. If *k* is high, CI has the potential to cause strong effects on the primary sex ratio, both because the baseline male production is by definition low under high *k*, and because there are many fertilized eggs available to be masculinized. For the female equation, the parameter *k* cancels out.

Appendix A gives explicit solutions for *Wolbachia* frequencies at the two non-trivial equilibria in these models. In Appendix B we explore the stability of the non-trivial equilibria and show numerically that when the two non-trivial equilibria exist, the lower frequency equilibrium is unstable and the higher one is stable (as claimed by Vavre et al., 2000). In Appendix A, we show analytically that the stable equilibrium frequency for *Wolbachia* infection exceeds one half when *f* ≤ 1, with the caveat that this applies to both sexes in the female-killing model, but only to the *Wolbachia* frequency in female hosts in the masculinization model. Regardless of this caveat, numerical examples show that the ability of *f >* 1 to create stable equilibria below one half for both sexes extends to both models (for examples of equilibria, see Fig. 5). The proofs in Appendix A follow the same logic as for the diplodiploid model: the unstable and stable equilibria are symmetric around a point *r* (for each sex), which is never below one half if *f* ≤ 1. Hence, again, a stable equilibrium close to one half may be vulnerable to the population falling under the invasion barrier if stochasticity causes fluctuations in *Wolbachia* frequency, as discussed above with the diplodiploid model.

**Figure 5.**
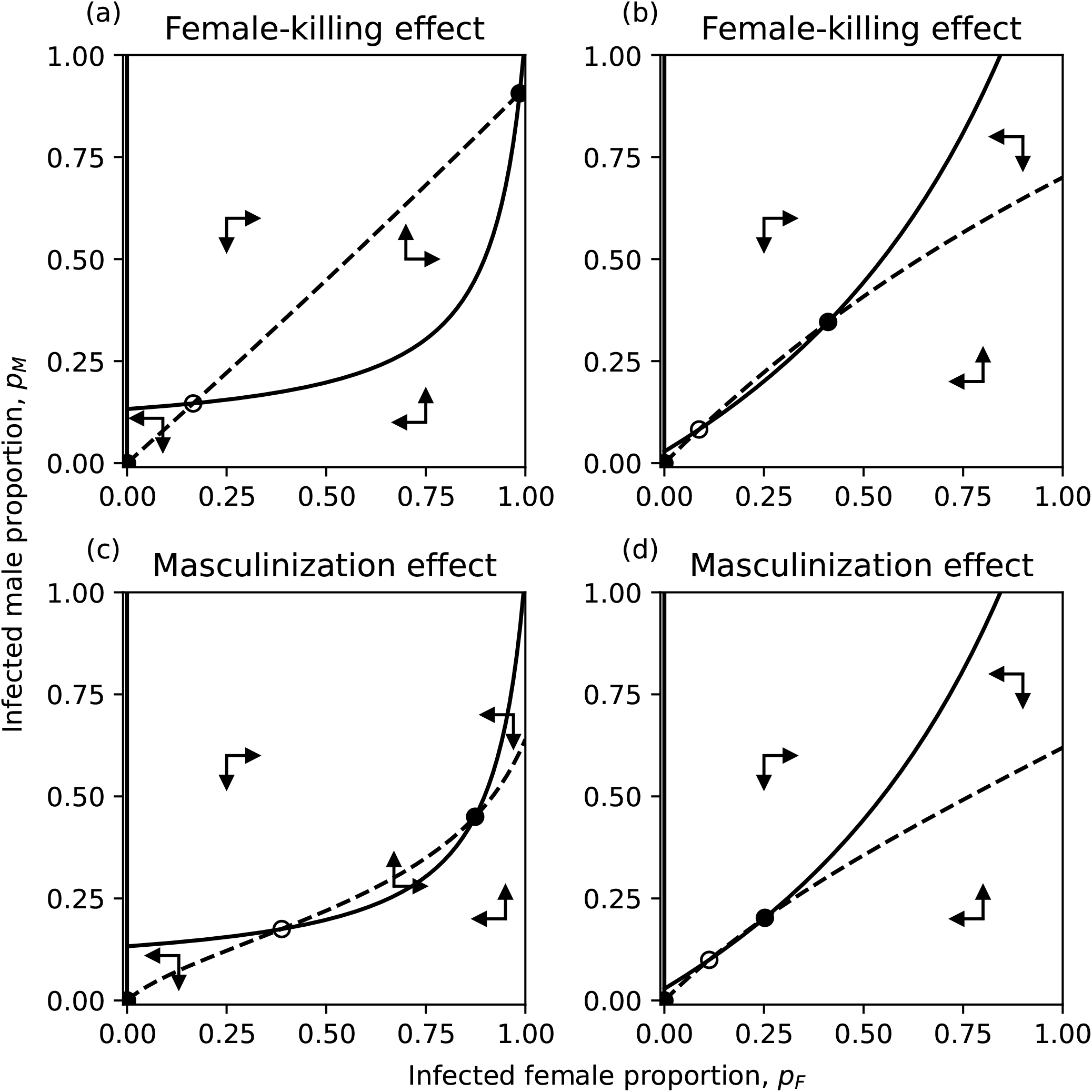
Equilibria of the two haplodiploid models in (*p*_*F*_, *p*_*M*_)-space. In left column *f <* 1, right *f >* 1. Dot: stable equilibria, circle: unstable equilibrium. Dashed line: male null-cline (Δ*p*_*M*_ = 0), solid line: female null-cline (Δ*p*_*F*_ = 0). a) The stable equilibrium is close to fixation and invasion threshold exists. b) Low frequency stable equilibrium with invasion threshold. c) The non-trivial stable equilibrium at 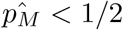, although *f <* 1. d) Low frequency stable equilibrium with invasion threshold. Parameters: a) and c) *f* = 0.95, *t* = 0.92, *L* = 0.95, *k* = 0.9 (only c); b) and d) *f* = 1.4, *t* = 0.7, *L* = 0.7, *k* = 0.5 (only d).

The infection frequency in males generally does not have to be the same as in females, because of the potential for additional female mortality and of the two different ways in which males can be produced: the egg not being fertilized in the first place, or via masculinization. In Appendix C, were prove analytically that infection frequency in females is always higher than or equal to that in males, for both the female-killing model and the masculinization model.

The additional flexibility actually brings about the possibility, in the masculinization case, that the frequency of infection in males can stabilize below one half even with *f* = 0.95 *<* 1 (Fig. 5c). The parameter range for this type of situation is narrow whenever *f* is low (Fig. 6), though. In addition, the female infection frequency was very high whenever *f* ≤ 1, so that the average infection frequency was above one half even in the few cases where male infection frequency was below one half under the masculinization model (Fig. 7). Such cases were found with high fertilization rate (*k*) yielding a strongly female-biased sex ratio, hence the average infection frequency was close to that found in females. Overall, low infection frequencies seem very rare unless *f >* 1.

**Figure 6.**
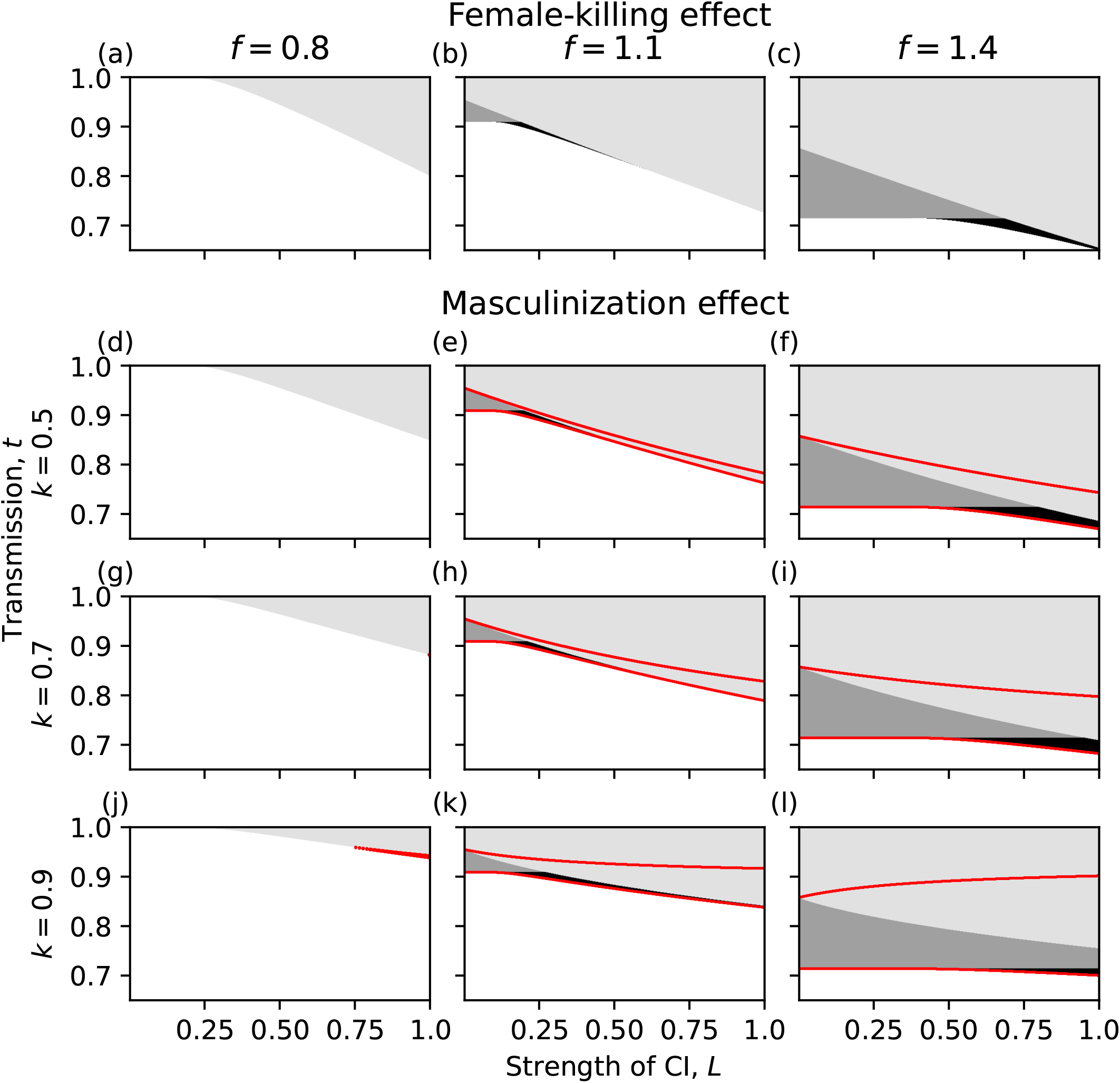
Classification of equilibria of the haplodiploid models across parameter space. Columns from left to right with *f* = 0.8, *f* = 1.1 and *f* = 1.4. First row: female-killing effect; other rows: masculinization effect model with different values of *k*. White: only the trivial equilibrium at zero. Light grey: non-trivial stable equilibrium with 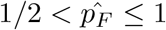. Dark grey: 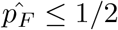 without invasion threshold. Black: 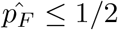 with invasion threshold. The red lines outline two thresholds, between which 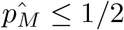. Note that the vertical *t*-axes span only high values, where non-trivial equilibria exist.

**Figure 7.**
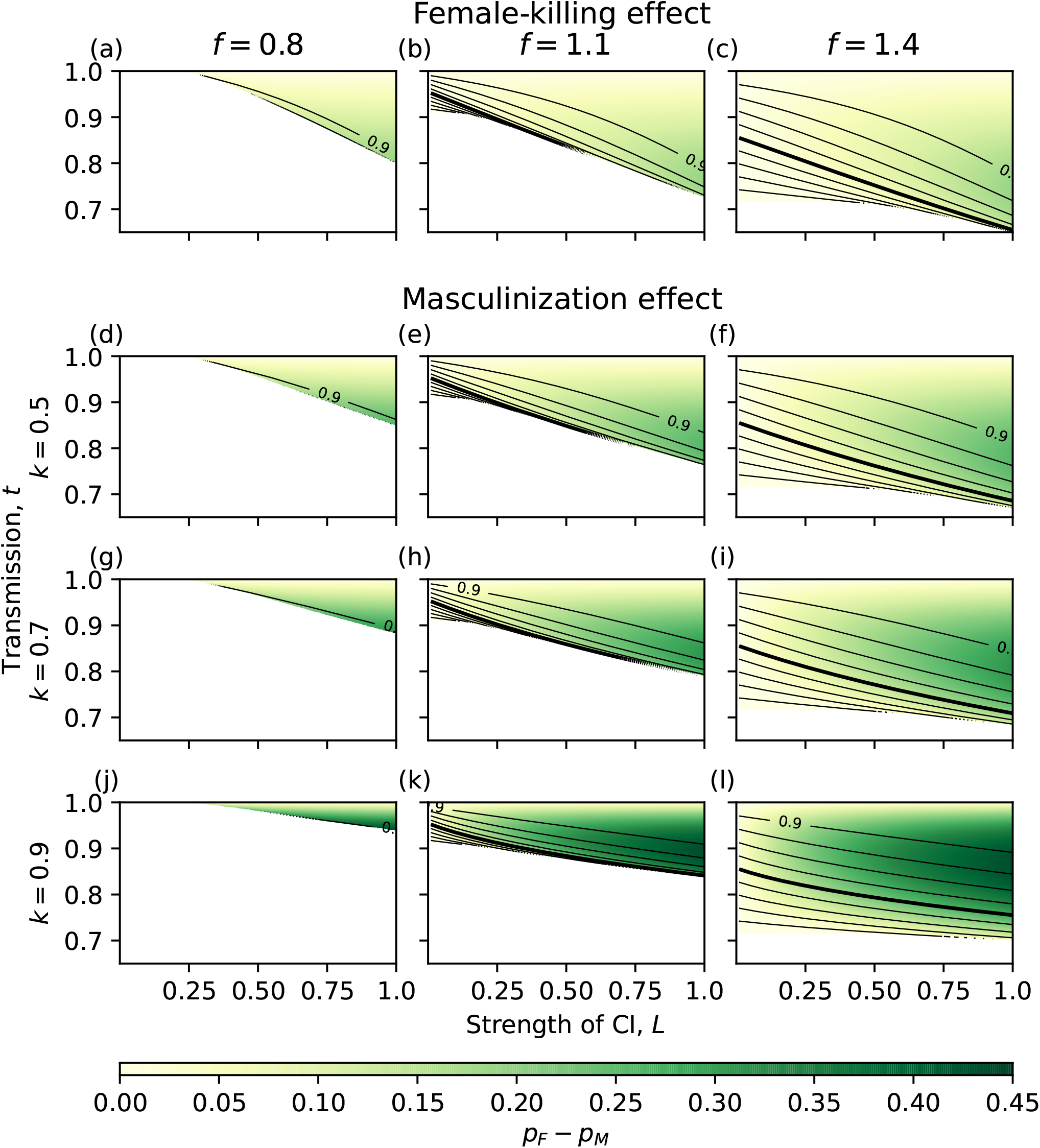
Equilibrium infection frequencies of the haplodiploid models across parameter space. Columns from left to right with *f* = 0.8, *f* = 1.1 and *f* = 1.4. First row: female-killing effect; other rows: masculinization effect model with different values of *k*. Black contour lines show infection frequency in females (with gaps of 0.1), while the color shows difference between female and male infection frequency according to the colorbar in the bottom. Thick solid contour marks one half frequency. Females always have higher infection frequency than males.

## Discussion

The traditional model of the dynamics of symbionts inducing cytoplasmic incompatibility predicts stable infection frequencies above one half in diplodiploid systems (Turelli, 1994; Turelli and Hoffmann, 1995). However, the prediction of high stable frequencies is tightly linked to an assumption of direct fitness effects of infection being either neutral or detrimental. Our findings, that low-frequency (below one half) stable equilibria can exist, are as such not new: examples were presented by Zug and Hammerstein (2018), but these authors did not elaborate on when exactly low frequencies can be expected to prevail. We show analytically that they can occur only if the relative fecundity of the infected individuals is higher than one.

The association between negative fitness effects and high frequencies is explained by positive frequency-dependence. CI penalizes uninfected individuals’ reproduction, but cannot do so very efficiently if infections are rare; therefore, infections must overcome an invasion barrier first but can thereafter spread to high frequencies (only counteracted by leaky transmission). Positive fitness effects remove the difficulties of spreading from very low frequencies, and as a whole this creates conditions for stable equilibria at low frequency. Similar results hold for haplodiploid systems, with the exception of male infection frequency in masculinization-type CI, which can stay below one half even with relative fecundity less than or equal to one.

The simplest case of low stable frequencies occurs when the effective fitness of rare infected individuals is higher than that of uninfected individuals (*ft >* 1). In that case, a rare infection can spread as there is no invasion barrier (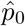 is unstable; see also Zug and Hammerstein, 2018), and the infection frequency rises to levels dictated by the strength of CI, efficiency of transmission, and relative fecundity. With very weak CI and leaky transmission, the stable equilibrium can be very close to zero (e.g. 0.06 in Fig. 2c, d). A more complex situation occurs when the relative fecundity of infected individuals exceeds one but not by much, so that relative fecundity above one combines with effective fecundity below one (*f >* 1 with *ft <* 1). In that case, the invasion threshold exists, but it is still possible to have a stable equilibrium below one half, a finding that has not been demonstrated before. As the previous analyses usually assumed equal or lower fecundity of infected individuals, they consequently observed stable equilibria above one half.

Note that similar results arise when considering a sexual population containing a lineage with a wild-type symbiotic strain and a lineage with a fitness-increasing symbiotic strain with non-perfect transmission. As shown by Zug and Hammerstein (2018), the condition *ft >* 1 is necessary and sufficient for the successful invasion of the fitness-increasing strain, as it provides the strain with higher effective fecundity than the wild-type lineage. They also mentioned this being analogous to the mutation-selection balance of haploids (Hoffmann and Turelli, 1997). Along the same lines, when studying the possibility of sex-ratio determination of CI-inducing symbionts, Egas et al. (2002) conclude that the symbiont can invade “if the proportion [of] infected daughters produced by infected females is bigger than the proportion [of] daughters produced by uninfected females (when mated with uninfected males)”, again returning to higher effective fecundity of the infected individuals.

Given that the consequences of higher relative fecundity are not only clear but also, in the form of examples, present in earlier analyses, why have positive fitness effects not been discussed explicitly before? A typical discussion of *Wolbachia* focuses on the fitness deficit caused by infection (Hoffmann, Turelli, and Harshman, 1990), which creates an interesting paradox: why should a symbiont with negative fitness effects on its host be so common in natural populations? The discovery of reproductive manipulation (such as CI) solved this conundrum. The possibility of positive fitness effects has thereafter not attracted much attention, despite the relevant equations remaining valid when *f >* 1. As an exception, Zug and Hammerstein (2018) recently explored positive fitness effects, with a focus on the evolution of the infectious agent and comparison of different symbionts (non-manipulating, CI, male-killing).

Besides showing that the stable equilibrium almost always exceeds one half when *f* ≤ 1, we also argue that infection frequencies slightly above one half are not expected to be robust in nature. Our model analysis thus suggests that positive fitness effects of CI-inducing *Wolbachia* infection are expected to be found in those natural populations that show low and stable densities of infection over time. Stable infection frequencies below one half and even slightly above it are likely paired with positive fitness effect unless another dynamic can explain the situation. Besides the fitness benefit demonstrated here, low frequencies may of course be observable during ongoing invasion or result from source-sink dynamics (e.g. Flor et al., 2007; Telschow et al., 2007).

It would be ideal to complement our findings with concrete empirical examples of stable CI-inducing *Wolbachia* strains that are maintained at low frequencies in their host populations. Unfortunately, it is difficult to highlight one specific case that perfectly illustrates our findings. Although it is well appreciated that *Wolbachia* is common among species while not necessarily reaching high frequencies within a species (Sazama et al., 2019), studies typically do not provide much information regarding additional fitness effects when populations have been screened for *Wolbachia* infection (Russell et al., 2012). However, there is some empirical evidence for *Wolbachia* strains that elevate host fitness, where the benefit may manifest itself, for example, as higher fecundity (Fry et al., 2004) or protection from viral diseases (Chrostek et al., 2013), and also evidence of evolution towards mutualism (higher fecundity) in natural *Wolbachia* populations (Weeks et al., 2007). Often, there is no information on whether the symbiont induces CI or any other phenotype in their hosts (Duplouy, Couchoux, et al., 2015). The stability properties of reported low frequencies are rarely evaluated, as most studies report prevalence of the symbiont at a single time point; recurrent screenings of the same natural populations over the course of several generations are rare (however, see Duplouy, Couchoux, et al., 2015; Duplouy, Nair, et al., 2021; Kriesner et al., 2013). Hence, the changes in the infection frequencies through generations remain unknown, and it is difficult to determine stability of the infection frequency, let alone any existence of invasion barriers or other dynamical features of the system.

Another testable prediction from the presented models is that in the haplodiploid models the female infection frequency seems to be consistently higher than the male infection frequency. Additionally, the masculinization type seems to predict lower equilibrium frequencies than the female-killing type. The latter observation might be explained, at least partially, by the fact that masculinization allows for excess production of uninfected males (compared to the zygotes dying), which “dilutes” the infection.

Further possibility for extension of this work would be to study the total size of the host population and the effect of *Wolbachia* on that. Also, understanding the details of dynamics approaching to fixed points would shed light on the diversity of possibilities that could be observed in nature. Both the host population size and the dynamics before reaching a stable equilibrium would be of interest when applying *Wolbachia* as a control agent.

Our model highlights that when strains are found at low frequencies in their host populations, there are additional possibilities to them being either in the process of being eliminated or of invading (Duplouy, GD Hurst, et al., 2010; Kriesner et al., 2013) their host population: a low frequency can be a stable outcome. As the mitochondrial haplotype will hitchhike a succesfully spreading maternally inherited *Wolbachia* infection, the state of potentially ongoing change can partially be answered with the study of the associated mitochondrial haplotype diversity in the host population, but often even this data is not available (but see Duplouy, GD Hurst, et al., 2010; Richardson et al., 2012). Long-term surveys of host populations and their infection status through time, as well as the study of their ecology are therefore needed. The recent clear results from the effective control strategy against human-born disease such as Dengue (suppressing its vector, *Aedes aegypti*) have clearly shown that long-term surveys are possible and informative for endosymbionts (e.g. Ryan et al., 2019).

## Acknowledgements

We are thankful for helpful comments of Roman Zug and Martin Kapun on the earlier versions of this work. We are grateful for exceptionally thorough and constructive feedback of three anonymous reviewers and the editor Jorge Peña. Preprint version 5 of this article has been peer-reviewed and recommended by Peer Community In Ecology (https://doi.org/10.24072/pci.ecology.100104).

## Fundings

Funding was provided by the Swiss National Foundation to HK, PK and CdV. CdV was also supported by an Academy of Finland grant (no. 340130, awarded to Jussi Lehtonen). AD was funded by Academy of Finland grant number 321543.

## Conflict of interest disclosure

The authors declare that they comply with the PCI rule of having no financial conflicts of interest in relation to the content of the article. The authors declare the following non-financial conflict of interest: AD is a recommender for PCI Ecology.

## Data, script, code, and supplementary information availability

Matlab (v. R2019b) and Mathematica (v. 12.1.1.0) were used for solving and analysing equations. The images were created and numerical analysis conducted in Python (v. 3.7.4) using packages numpy (v. 1.21.5, Harris et al., 2020) and matplotlib (v. 3.4.2, Hunter, 2007). Source code is provided in the Zenodo repository: https://doi.org/10.5281/zenodo.6443870 (Karisto et al., 2022).

## Appendix A: Limits for CI frequency in haplodiploid systems

### Female-killing effect

When CI causes mortality of the female embryo in haplodiploid species, the population dynamics follow the model in Eq. (3). The non-trivial equilibria of the model are

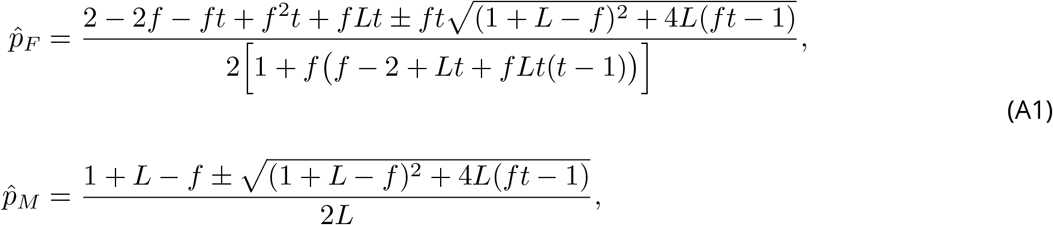

where pluses belong to the non-trivial stable equilibrium 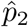 and minuses to the unstable equilibrium 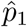 when they exist (Vavre et al., 2000). The stability of these is explored numerically in Appendix B.

Note that for the equilibrium 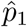 we can see that when *ft* = 1, 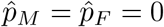 and additionally when *ft >* 1, 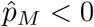, hence the “invasion threshold” coincides with 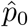 at *ft* = 1 and is not in the biologically relevant range when *ft >* 1, similar to the diplodiploid model.

We use the same technique as for diplodiploid model, showing that the “real part” *r* (the part that is necessarily real, i.e. excluding the square root part) of the equilibrium frequency is below one half if and only if *f >* 1, and hence 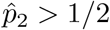 for *f* ≤ 1. For the male frequency, it is easy to see that

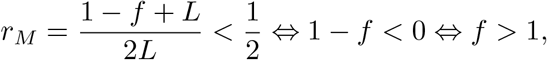

but for the female frequency the proof is somewhat more complicated. In that case

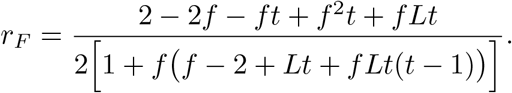

The denominator of *r*_*F*_ divided by two (i.e. the expression within square brackets) is a quadratic function of *f*,

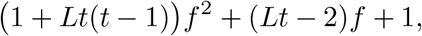

which is always positive, as the intercept is positive and no real roots exist (discriminant *Lt*2(*L*−4) *<* 0). As the denominator of *r*_*F*_ is always larger than zero, we can multiply the inequality *r*_*F*_ *<* 1*/*2 with the denominator, resulting in an equivalent inequality

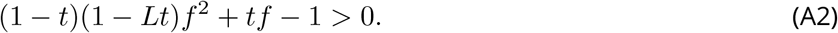

In Eq. (A2) the intercept of the left hand side is negative and the quadratic coefficient (1−*t*)(1−*Lt*) *>* 0. Hence, the equation holds true for positive *f* only when *f* is larger than the larger root, say *f*_lim_, of the quadratic function in the left hand side of Eq. (A2). Only if *f*_lim_ *<* 1, can *f* ≤ 1 fulfil the inequality in Eq. (A2) and equivalently *r*_*F*_ *<* 1*/*2. Let us now investigate in what conditions *f*_lim_ *<* 1:

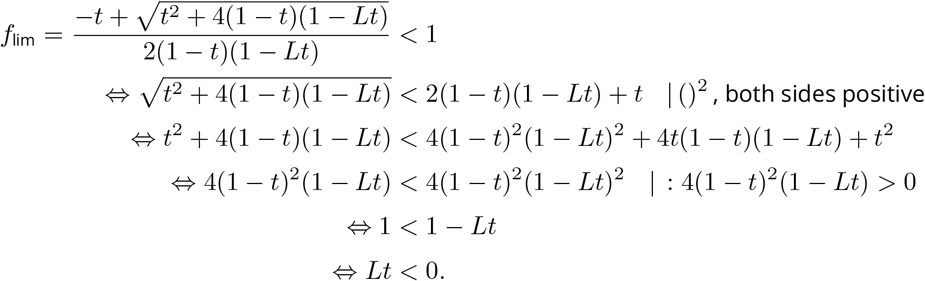

Thus, the lower limit *f*_lim_ is below one only if *Lt <* 0, which is never true. Hence the limit is always at least 1, which implies that *r*_*F*_ *<* 1*/*2 is true only if *f >* 1. This concludes the proof that 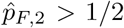 for the female-killing type CI when *f* ≤ 1.

### Masculinization effect

The CI can alternatively lead to deletion of the paternal genome, turning the diploid egg into a haploid that develops as male. In that case, the population dynamics follow the model in Eq. (4). The non-trivial equilibria of the model are

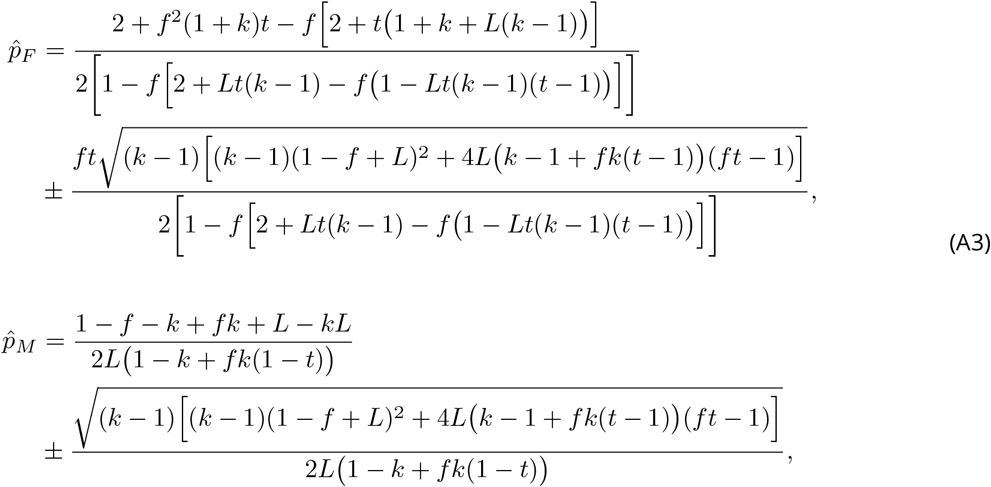

where pluses belong to the non-trivial stable equilibrium 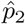 and minuses to the unstable equilibrium 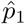 when they exist (Vavre et al., 2000). The stability of these is explored numerically in Appendix B.

We show that 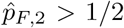 whenever *f* ≤ 1 using the same logic as before, i.e. we show by contradiction that for any *f* ≤ 1 and valid parameter values, “the real part” of the solution, *r*_*F*_, cannot be less than 1*/*2. Thus, *r*_*F*_ ≥ 1*/*2 and hence also 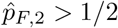 as it is larger than *r*_*F*_, which is

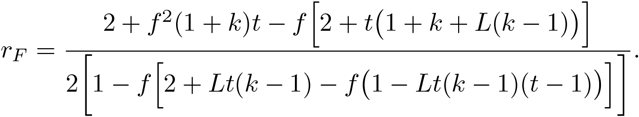

The denominator of *r*_*F*_ is a linear function of *k*:

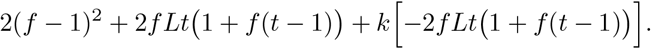

When *f* ≤ 1, the slope of that function is negative (−2*f Lt*(1 + *f* (*t* − 1)) *<* 0) and the root satisfies

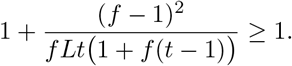

Thus, with *f* ≤ 1 the denominator is positive for *k <* 1, and the inequality *r*_*F*_ *<* 1*/*2 simplifies to the equivalent condition

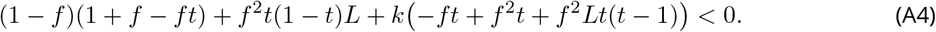

The left hand side is again a linear function of *k*, which has an intercept that is always positive for *L* ∈ (0, 1] as it is linear in *L* and has positive intercept and slope. The linear function of *k* in Eq. (A4) has one root at

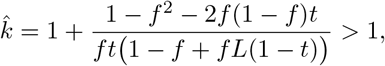

as the second term is positive for *f* ≤ 1. The denominator of the second term is obviously positive and the numerator is also positive, as it is linear in *t*, having negative slope and root (1 + *f*)*/*2*f* ≥ 1 when *f* ≤ 1.

Thus, Eq. (A4) has positive intercept and 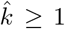 holds, which means that the left hand side of Eq. (A4) is positive for all *k* ∈ (0, 1[and hence the inequality in Eq. (A4) is never true for valid values of *k*. As Eq. (A4) is equivalent with *r*_*F*_ *<* 1*/*2, this implies that the real part of the female equilibrium frequency is never less than 1*/*2 with valid parameter values (assuming *f* ≤ 1). The actual stable equilibrium is even higher than *r*_*F*_, which concludes the proof that female frequency is never below 1*/*2 at the stable equilibrium.

Finally, for the infection frequency in males under the masculinization effect model, it is easy to prove by example that the frequency at the stable equilibrium indeed can be below 1/2 even with *f <* 1. In Fig. 5c, the parameters are *f* = 0.95, *t* = 0.92, *L* = 0.95, *k* = 0.9 while 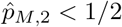.

In conclusion, the infection frequency in females follows the same pattern as female-killing type and diplodiploid systems with symmetry of the stable equilibrium and the unstable equilibrium around a point that is above one half for all *f* ≤ 1, but the infection frequency in males makes an exception as it can go below one half even with *f <* 1.

## Appendix B: Local stability analysis of the equilibria of CI models

In this appendix we will use linear stability analysis to calculate the local stability of the equilibria of the different models presented in the main text.

### Diplodiploid model

The dynamics of *Wolbachia* frequency in both males and females in this model is given by Eq. (1) in the main text, here repeated in slightly different form that specifies the time *T* explicitly:

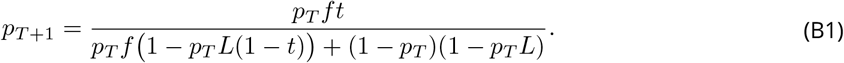

For notational convenience, define a function *g* as

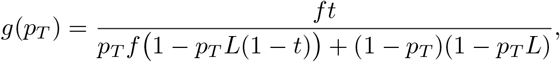

so that

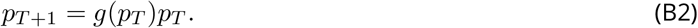

Equilibria of this discrete time dynamical system satisfy

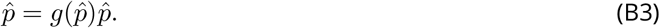

An equilibrium, 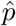, is locally stable if and only if the absolute value of the derivative of *p*_*T* +1_ with respect to *p*_*T*_ is smaller than one (see e.g. Otto and Day, 2007), that is,

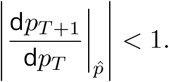

We will denote this derivative evaluated at equilibrium 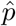 as 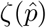.

Therefore to calculate the stability of the equilibria of our diplodiploid model, we need the derivative of *p*_*T*+1_ with respect to *p*_*T*_. Differentiating Eq. (B2) gives

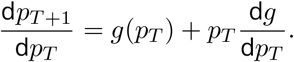

Next we want to evaluate this derivative at the equilibria. Equation (B1) has three equilibria, solved from equation *Ap*2 + *Bp* + *C* = 0, as shown in the main text and repeated here for convenience:

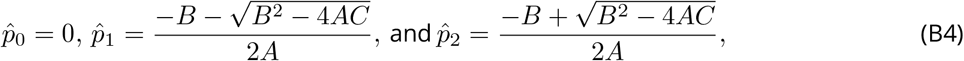

where *A* = *L*(1 − *f* (1 − *t*)), *B* = −1 + *f* − *L* and *C* = 1 − *ft*.

The stability of the first equilibrium, 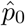, is given by

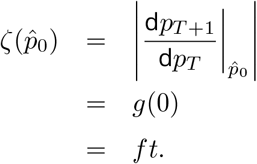

The equilibrium 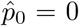 is therefore stable if and only if *ft <* 1. At *ft* = 1, the equilibria 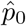 and 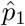 coincide, and when 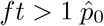 is unstable. Note that this has implications on the existence of invasion threshold, since there is no invasion threshold if 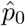 is unstable (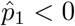 when *ft >* 1). In that case the infection frequency will increase from any initial frequency until it reaches a stable equilibrium.

The stability of the second equilibrium, 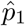, is given by

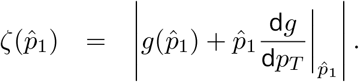

The first term 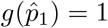 when 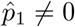, because of Eq. (B3). The second term is

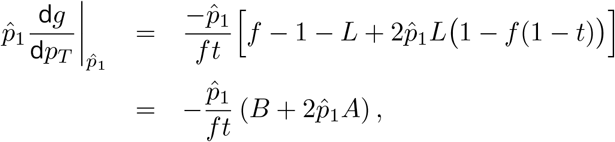

and hence

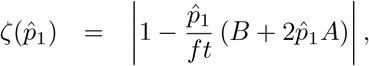

or using Eq. (B4),

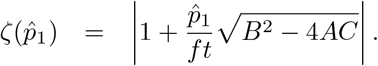

Stability requires that 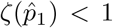. Since *ft >* 0 always holds, equilibrium 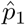 is unstable whenever 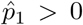. The first non-trivial equilibrium is therefore always unstable when it is positive, in agreement with Hoffmann, Turelli, and Harshman (1990).

Similarly, the stability of the second equilibrium, 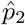 is given by

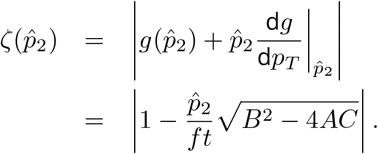

The equilibrium 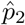 is therefore stable if and only if

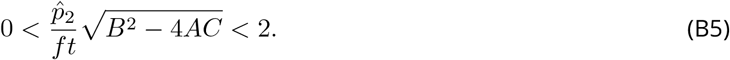

If 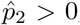, then the left-hand side of this inequality is true. To check the right-hand side of the inequality, we substitute the expression of 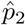 in Eq. (B4) into Eq. (B5) to obtain:

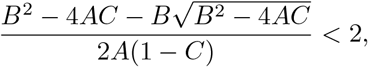

or equivalently

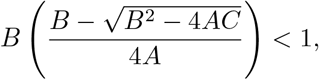

which can be written as

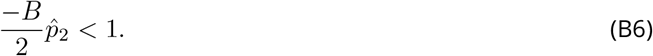

Now, if *B <* 0 (i.e. *f <* 1 + *L*) we can multiply both sides of inequality by −2*/B* to get an equivalent stability condition

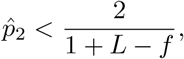

which is always true since 0 *<* 1 + *L* − *f <* 2 and hence the right-hand side is larger than 1. Therefore as long as 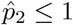 this condition is fulfilled and 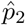 is a stable equilibrium. On the other hand, if *B* ≥ 0 (i.e. *f* ≥ 1 + *L*), the left hand side of Eq. (B6) is zero or negative since −*B <* 0 and the inequality is true for all 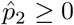. Hence, 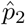 is always stable if it exists within the biologically relevant range between 0 and 1.

### Haplodiploid models

The *Wolbachia* models with haplodiploid sex determination in the main text are both frequency-dependent models that in general we can write as follows

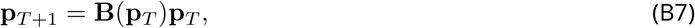

where **p**_*T*_ is a vector with the frequency of *Wolbachia* infection in females and males as its entries. Moreover, both the female-killing effect model, and the masculinization effect model have the following general shape:

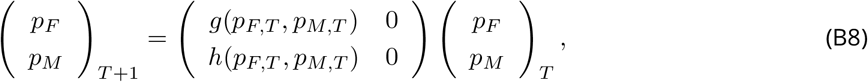

where the functions *g* and *h* are different for the female-killing model and the masculinization model, and they are presented in the sections below.

Equilibria, 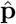, of this discrete time dynamical system satisfy

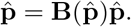

The stability of an equilibrium to small perturbations is determined by the absolute value of the dominant eigenvalue of the Jacobian matrix of the model evaluated at the equilibrium. The equilibrium is stable if the absolute value of the dominant eigenvalue of the Jacobian matrix is smaller than one. The Jacobian matrix is obtained by differentiating Eq. (B7) and when evaluated at an equilibrium it is

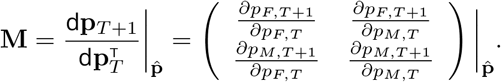

As both models have similar structure (Eq. B8) their Jacobian matrices have the general form

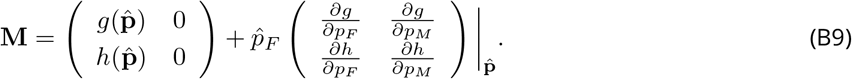

### Female-killing effect

We consider the female-killing effect first (Eq. (3) from the main text in recursion form with explicit time *T*):

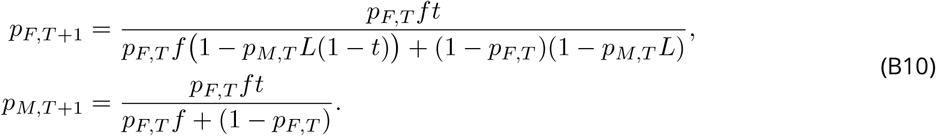

Therefore, if we define,

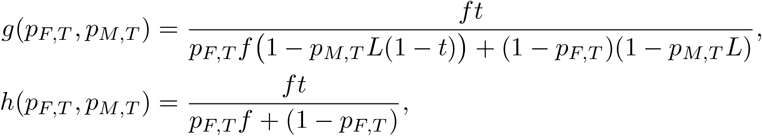

then we can write the recursion equations (B10) in the matrix form given above by equation (B8).

For the trivial equilibrium, 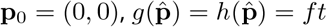 and we have

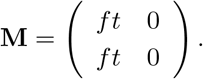

The eigenvalues of this Jacobian matrix are 0 with eigenvector (*p*_*F*_, *p*_*M*_) = (0, 1), and *ft* with eigenvector (*p*_*F*_, *p*_*M*_) = (1, 1). The zero eigenvalue is associated with perturbations in the directions of males only, which makes sense biologically: these perturbations decay back to the zero equilibrium since males do not transmit *Wolbachia*. The largest eigenvalue of the **p**_0_ = (0, 0) equilibrium is *ft*. Similarly to the diplodiploid model, the trivial equilibrium is therefore stable if *ft <* 1, and unstable if *ft >* 1.

The non-trivial equilibria of this model are given in Eq. (A1) and repeated below,

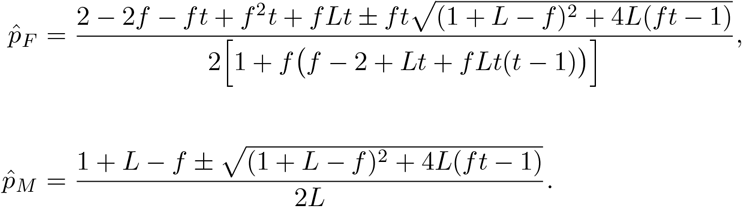

To calculate the Jacobian matrix for the non-trivial equilibria, we need to calculate the derivatives in Eq. (B9). It will be useful to note that for the non-trivial equilibria, 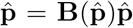 implies that 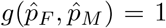, and 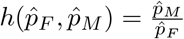. It follows that

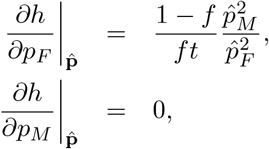

and

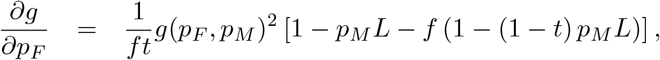

and therefore

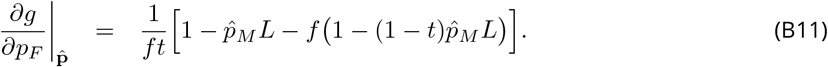

The equilibrium condition 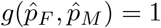 can be rewritten as

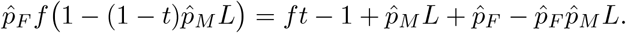

Multiplying Eq. (B11) with 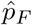 in Eq. (B9), and substituting the above expression yields

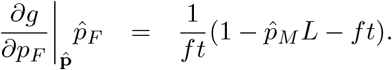

Similarly,

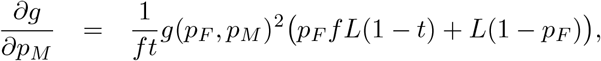

and

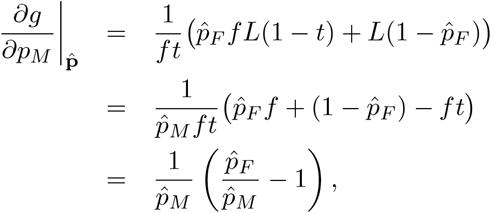

where we have used 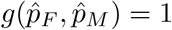 and 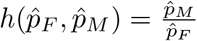 for the last two lines, respectively.

Putting all of this together, we get the following Jacobian matrix,

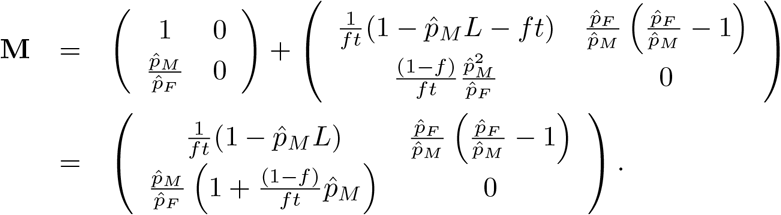

The eigenvalues of this Jacobian, denoted by *λ*, are given by the following quadratic equation

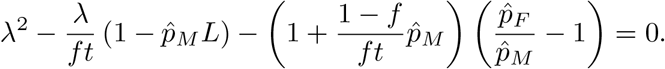

If the absolute value of the dominant eigenvalue of the Jacobian is below one, then the equilibrium is stable. The eigenvalues are given by

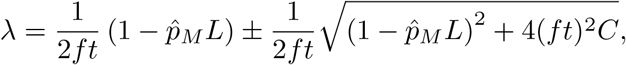

where

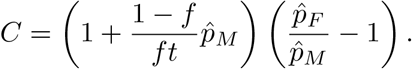

Since *L* ≤ 1, the first term of *λ* is positive and the eigenvalue with the largest absolute value is

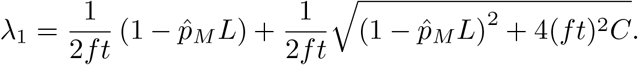

We could not determine stability of the equilibria analytically. Instead, we used Python to explore the parameter space *{*0.01 ≤ *L, t* ≤ 1*} × {*0.01 ≤ *f* ≤ 3*}* sampling values for each parameter uniformly with 0.01 interval, which resulted in 3 000 000 parameter combinations. Whenever an equilibrium (either 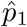 or 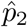) appeared in the biologically meaningful range 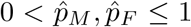, we analysed its stability. For all cases where |*λ*_1_| ≠ 1 (i.e. linear stability analysis did not fail) the linear stability analysis concluded that 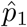 was unstable and 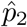 was stable.

### Masculinization effect

For the masculinization effect, the dynamics for females and males, respectively, are given by Eq. (4) in the main text (here in recursion form):

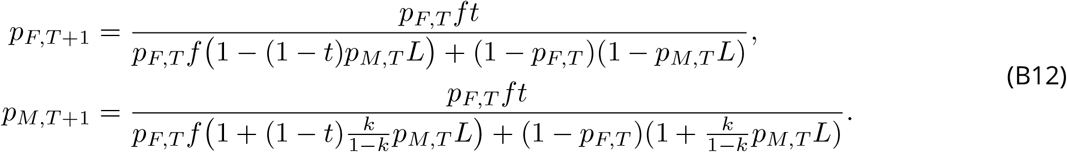

Therefore, if we define,

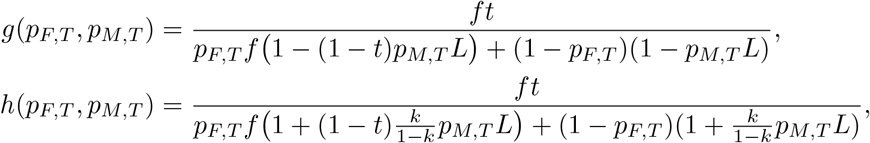

then we can write the recursion equations (B12) in the matrix form given above by equation (B8).

The non-trivial equilibria of this model are given by Eq. (A3) and repeated here:

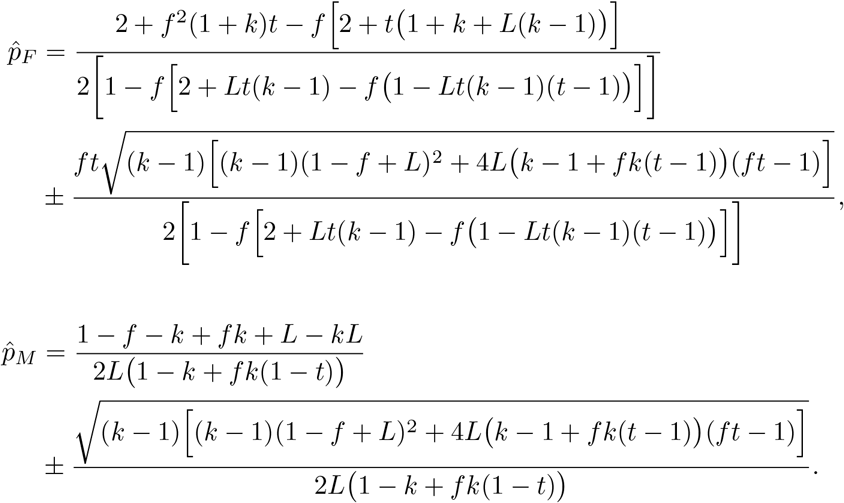

As before, at the trivial equilibrium, 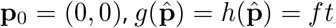, and therefore

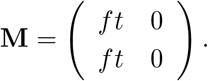

Again the eigenvalues of this Jacobian matrix are 0 with eigenvector (*p*_*F*_, *p*_*M*_) = (0, 1), and *ft* with eigenvector (*p*_*F*_, *p*_*M*_) = (1, 1). The zero eigenvalue is associated with perturbations in the directions of males only. The largest eigenvalue of the **p**_0_ = (0, 0) equilibrium is *ft*. Similarly to the diplodiploid model, the trivial equilibrium is therefore stable if *ft <* 1, and unstable if *ft >* 1.

For the non-trivial equilibrium, we need to calculate the derivatives in the formula for the Jacobian again (Eq. (B9)):

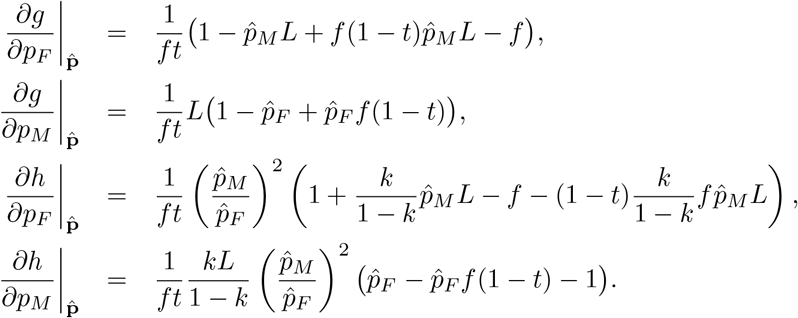

Using 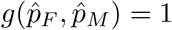 and 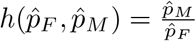, we can simplify

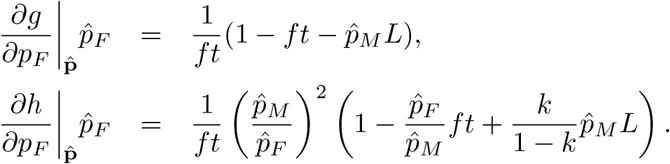

Putting it all together gives the following expression for the Jacobian matrix,

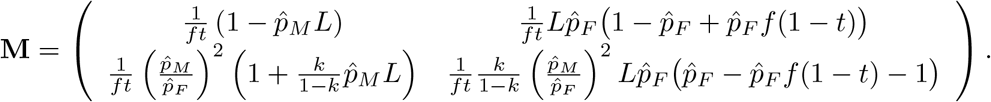

The eigenvalues of this Jacobian are given by the solution to

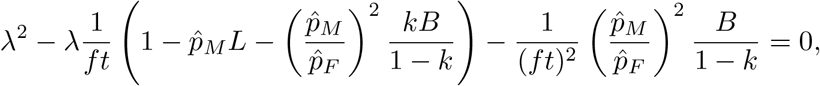

where

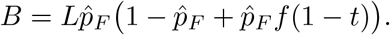

The eigenvalues are therefore equal to

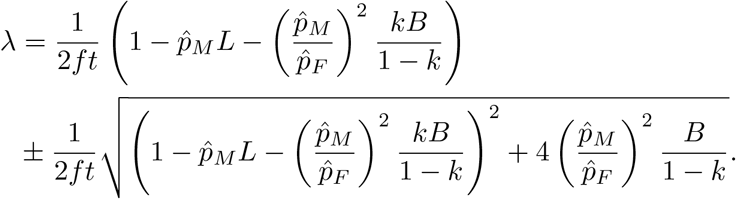

As with the female-killing model, we could not determine the stability of the equilibria analytically. Instead, we conducted similar numerical analysis in Python with 100 values for *{*0.01 ≤ *L, t* ≤ 1*}*, 300 values for *{*0.01 ≤ *f* ≤ 3*}*, and three values of *k* ∈ *{*0.5, 0.7, 0.9*}*, resulting in 9 000 000 parameter combinations in total. We calculated the values of the equilibria and determined their stability based on the larger absolute value of the eigenvalues above, whenever 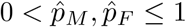. Again, the equilibria were unstable 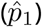 and stable 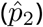 as expected, whenever they occurred within the biologically meaningful range and the leading eigenvalue had absolute value different from 1.

## Appendix C: *Wolbachia* frequency is higher in females than in males in the haplodiploid models

Here we show that in the haplodiploid models presented in the manuscript, the infection frequency in females is never lower than infection frequency in males.

### Female killing effect

We start from from the remark that 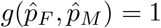. Hence,

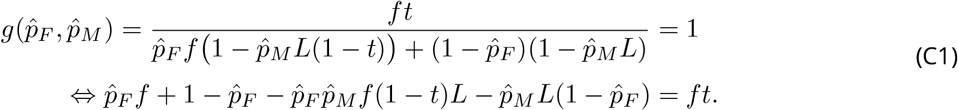

Next, we use 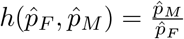, namely

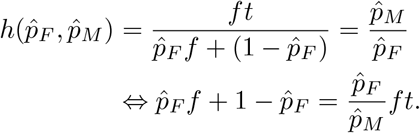

Substituting the latter row into Eq. (C1) yields

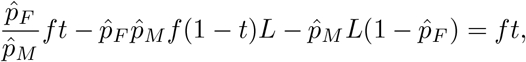

and therefore

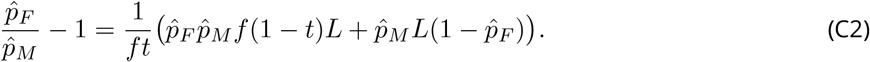

The right-hand side of Eq. (C2) cannot be negative with valid parameter values (including *f >* 1). Therefore 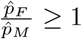, which means that the infection frequency in females is always higher than or equal to male infection frequency.

### Masculinization effect

We start again from 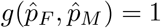, and 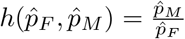. The first equality gives

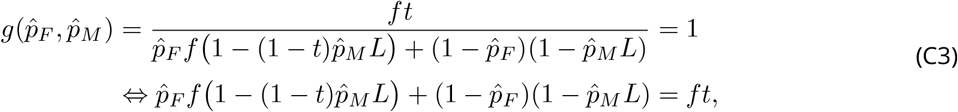

and the second gives

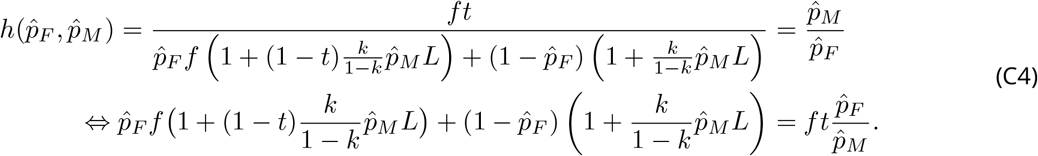

Subtracting the last row of Eq. (C3) from the last row of Eq. (C4) yields

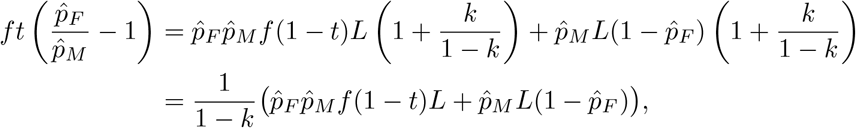

and therefore

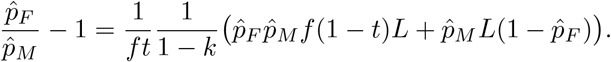

The right-hand side above is never below zero for any valid parameter values, and therefore 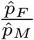 is never less than one. Hence, the female infection frequency is always higher than or equal to the male infection frequency.

## Notes

https://doi.org/10.5281/zenodo.6443870

### Competing Interest Statement

The authors have declared no competing interest.

### Summary of Updates

Improved formatting, added links to references and DOIs into reference list.

